# Highly diverse and unknown viruses may enhance Antarctic endoliths’ adaptability

**DOI:** 10.1101/2022.12.02.518905

**Authors:** Cassandra L. Ettinger, Morgan Saunders, Laura Selbmann, Manuel Delgado-Baquerizo, Claudio Donati, Davide Albanese, Simon Roux, Susannah Tringe, Christa Pennacchio, Tijana G. del Rio, Jason E. Stajich, Claudia Coleine

## Abstract

Rock-dwelling microorganisms are key players in ecosystem functioning of Antarctic ice free-areas. Yet, little is known about their diversity and ecology. Here, we performed metagenomic analyses on rocks from across Antarctica comprising >75,000 viral operational taxonomic units (vOTUS). We found largely undescribed, highly diverse and spatially structured virus communities potentially influencing bacterial adaptation and biogeochemistry. This catalog lays the foundation for expanding knowledge of the virosphere in extreme environments.

## Main

Viruses are among the most prevalent entities on our planet, with the ability to infect organisms across all Domains^1^. Sequencing advances are reshaping understanding of viral diversity across Earth’s diverse ecosystems, leading to a remarkable expansion of viral catalogs^2–4^. It is becoming clear that viruses play key roles in global biogeochemical cycles through the modulation of host population dynamics, and that the better-studied pathogenic viruses represent only a small fraction of the virosphere^5–7^. Further, through auxiliary metabolic genes (AMGs), some viruses can directly impact host metabolism to improve fitness^8^, including in extreme ecosystems.

Antarctic ice-free areas include several of the most inhospitable regions on Earth, among which is the Mars counterpart: the McMurdo Dry Valleys. In these locations, where rocks represent the main substratum, active life is possible for only a few specialized microorganisms; they survive by dwelling in porous rocks, forming self-sustaining ecosystems called endolithic communities^9,10^are the primary life-forms present assuring the balance and functionality of these otherwise inert ecosystems. Recent studies have shed light on their biodiversity and adaptation, particularly the evolution of new and peculiar taxa spanning bacteria, fungi and archaea^10–13^. However, the ecology and distribution of viral diversity from these communities remain wholly unknown and, to date, viral studies have instead focused on Antarctic freshwater lakes^14–16^, surrounding oceans^17–19^, and soils^20–23^.

Here, we provide a large-scale viral catalog from 191 Antarctic endolith metagenomes. We sampled 37 localities across a broad range of environmental (e.g. 4 rock typologies, different altitudes and sun exposure) and spatial conditions (i.e. Antarctic Peninsula, Northern Victoria Land, and McMurdo Dry Valleys) (Table S1; Fig. 2A). We aimed to (i) untangle viral diversity in these communities, (ii) predict AMGs and how they may drive the fitness of their hosts, and (iii) explore ecological patterns (e.g., biogeography). This catalog is the first step toward understanding the role of viruses in regulating biogeochemical cycling in the coldest and driest region on Earth. This information is also critical for elucidating the role of viruses in whole community adaptation in a scenario of global warming and expanding desertification^24^.

**Figure 1.**
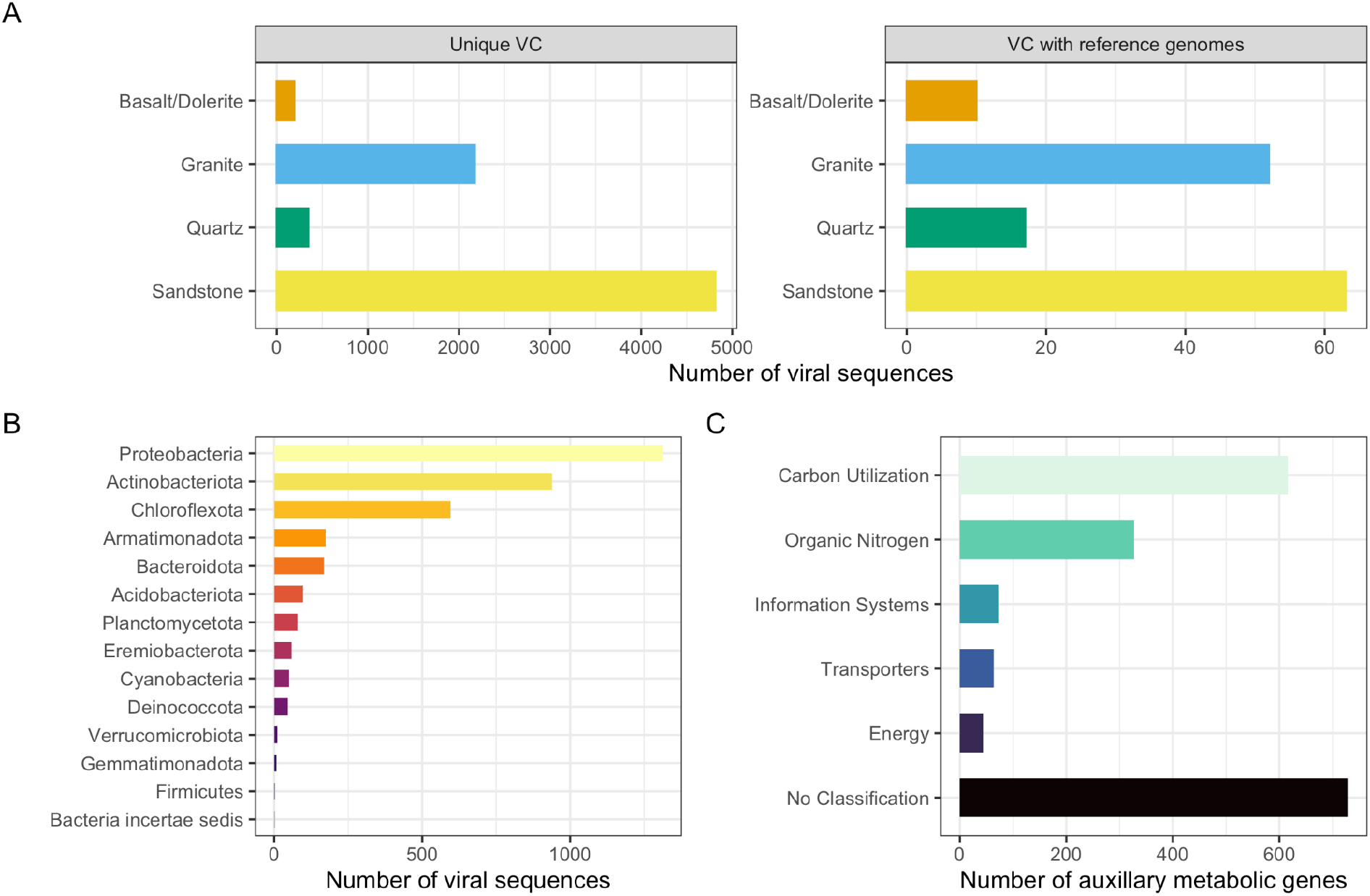
Antarctica is an underappreciated source of phage novelty. (A) Bar charts displaying the number of viral sequences placed in VCs colored by rock type and divided by whether the VC is clustered with reference genomes. (B) Bar chart displaying host predictions colored by predicted host phylum. (C) Bar chart showing the number of predicted phage AMGs summarized by DRAM-v distilled metabolic categories.

**Figure 2.**
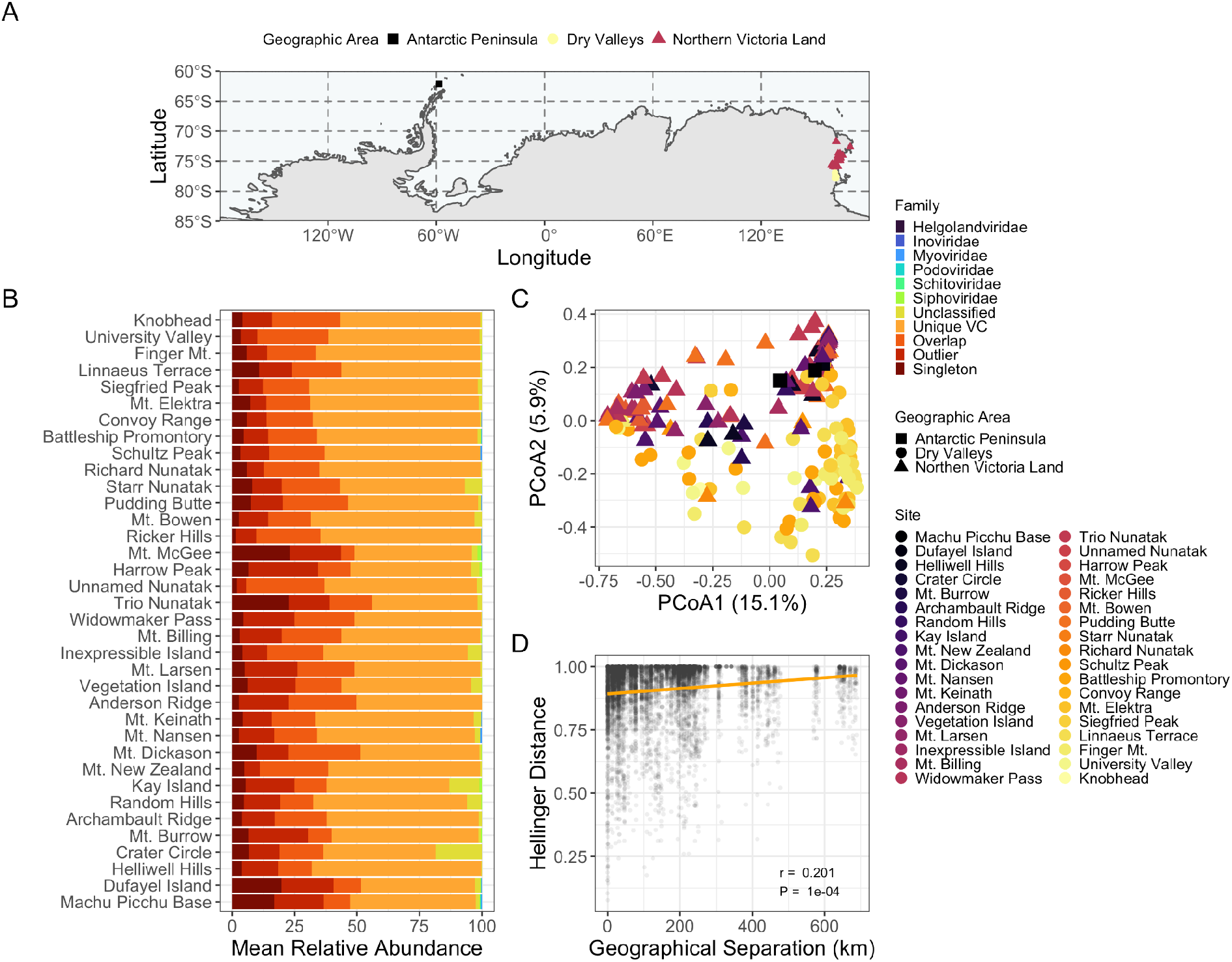
Spatial structuring of viral communities in Antarctic rocks. (A) Map showing collection sites with shapes and colors representative of the broad geographic area. (B) Stacked bar charts displaying the mean relative abundance of phage vOTUs at each site colored by predicted viral families. Sequences that were clustered into VCs with reference data are labeled by their taxonomy, sequences clustered without reference genomes are labeled “Unique VC”, while the rest are labeled based on their VContact2 status (i.e., singleton [share few or no genes with other genomes], overlap [share genes with genomes in multiple VCs], or outlier [share genes, but cannot confidently be placed in a VC]). (C) Principal-coordinate analysis (PCoA) visualization of Hellinger distances of viral communities. Samples are colored by site, with sites ordered by latitude, and have shapes based on geographic areas. (D) A scatter plot depicting a significant positive distance-decay relationship between sandstone viral community beta diversity (Hellinger distance) and geographical distance (km) between sites.

Using VirSorter2^25^, we predicted 101,085 viral sequences. We clustered these at 95% average nucleotide identity into 76,984 viral operations taxonomic units (vOTUS)^26^; we further used VContact2^27^ with INPHARED^28^ reference genomes to cluster phage vOTUs into 7,598 viral clusters (VCs), which approximate genus-level groupings based on gene-sharing networks. To keep analysis focused on the most robust catalog, we filtered this collection using community thresholds for length, detection, and quality (See Online Methods)^29–31^. The final viral catalog represented 14,797 viral sequences, including 2,695 prophage, which clustered into 11,806 vOTUs, of which 5,743 phage vOTUs (7,309 sequences) were successfully placed in 2,286 VCs; the final catalog was predicted to predominantly be dsDNA phage, though 15.2% of vOTUs may represent eukaryotic viruses (i.e. NCLDVs).

Our findings indicate that Antarctic rock communities host highly diverse and novel phage populations, with only 1.8% (41 out of 2,286) of the VCs including reference sequences. The remaining 98.2% were unique VCs (i.e., did not include reference genomes), and could represent novel phage genera, greatly expanding the known diversity of viruses. Of the 41 VCs that did include reference genomes, the majority were assigned to the *Caudoviricetes* class (formerly *Caudovirales* order) of tailed double-stranded DNA bacteriophage (Fig. S1). Many genomes have not yet been reclassified, leaving viral taxonomy in flux; under the new schema, most of the 41 VCs are unclassified^32^. The majority of unique VCs are represented by viral sequences from sandstone communities (Fig. 1A), which represents an optimum substratum, in terms of rock traits (e.g. porosity), for endolithic colonization^33^, but is also the most represented substratum in this work.

We further established host–virus linkages using NCBI BLAST against complete bacteria and archaea genomes from RefSeq, and Antarctic endolithic bacterial and archaeal metagenome-assembled genomes (MAGs) (see Online Methods)^34–36^ to explore the potential effects of viruses on host fitness, such as host-cell reprogramming through AMGs^37^. While we were unable to predict hosts for the majority of vOTUS, we observed that Proteobacteria, Actinobacteriota, and Chloroflexota were the most commonly predicted host phyla (Fig. 1B), which are thought to be core members of these communities^11,38,39^. Using predictions against the Antarctic MAGs, we predicted hosts for an additional 16.5% of viral sequences (Fig. S2).

We then sought to improve understanding on the functional profiles of retrieved phages using DRAM-v^40^. Notably, this catalog, which comprises metabolic novelty (39.3% of DRAM-v predicted AMGs had no distilled classification), may complement other available resources, which have largely been limited to coverage of human-related microbiomes (e.g. Li et al.^41^). Within identified functions, we found putative phage AMGs related to carbon, energy and nitrogen metabolisms (Fig. 1C). Specifically, within carbohydrate metabolism, glycoside hydrolases, glycosyltransferases, and carbohydrate-binding domains predominated. Within nitrogen metabolism, methionine degradation was the most prevalent module, and within energy, the dominant modules were related to electron transport and photosynthesis. This highlights the utility to connect vOTUs to Antarctic MAGs^11^ and to implement complementary techniques (e.g. single-cell genomics) to provide a deeper understanding of virus-bacteria dynamics. More importantly, these findings underscore the complexity of virus-driven element biogeochemical cycles in the rocks of Antarctica, which have traditionally been considered devoid of life.

Given the geographic spread of sampling (see Online Methods and Table S1; Fig. 2A), we assessed whether this catalog could be useful to answer ecological questions related to viral community dynamics. While the dominant vOTUs at each site were taxonomically unclassified and largely members of unique VCs and thus possible novel genera (Fig. 2B), when investigating between-sample diversity (beta diversity) we observed a significant pattern related to site specificity (Fig. 2C; PERMANOVA, *p* < 0.001), and further detected significant distance decay across sandstone communities (Fig. 2D; *r* = 0.197, *p* < 0.001), indicating clear latitudinal spatial structuring of viral communities. In further support of this, we were able to detect only 41.0% of vOTUs at more than one site, with 29.4% of vOTUs detected across two or more geographic regions and only 1.45% detected across all regions. Of the vOTUs detected across all regions, the majority were in unique VCs (66.7%) and none were in VCs with reference data. We hypothesize that this viral spatial structuring reflects the reported dispersion limitation and local composition and adaptation of hosts in these communities^11,12^. Similar spatial structuring has also been observed in grassland soil viromes, purportedly as a result of local assembly dynamics^42,43^.

This study represents the most exhaustive geographic endeavor to date to capture the viral genomic diversity across ice-free regions of Antarctica and the first large-scale effort to explore the virosphere in endolithic communities. This catalog is a comprehensive repository for exploring the diversity, function, spatial ecology, and host-virus dynamics of this enigmatic continent. We also unveiled a possible influence of some viruses on carbon, energy and nitrogen metabolisms under conditions of oligotrophy up to the limit for life sustainability; this may be a key role for the resilience of these communities. This work is a good model for exploring adaptability of microbial communities in a scenario of global warming and expanding desertification.

## Methods

### Study area

191 rocks colonized by endolithic communities were collected in thirty-eight sites in Antarctica including Antarctic Peninsula (*n* = 3), McMurdo Dry Valleys, Southern Victoria Land (n = 80), and Northern Victoria Land (*n* = 108) during more than 20 years of Italian Antarctic Expeditions. Different rock typologies (sandstone *n* = 141, granite *n* = 43, quartz *n* = 5, and basalt/dolerite *n* = 2) were sampled. Samples were collected along a latitudinal transect ranging from -62.10008 -58.51664 to -77.874 160.739 at different environmental conditions namely sun exposure (northern sun exposed and southern shady rocks) and an altitudinal transect from sea level to 3,100 m above sea level (a.s.l.) to provide a comprehensive overview of Antarctic endolithic diversity (Supplementary Table 1). The presence of endolithic colonization was assessed by direct observation in situ. Rocks were excised using a geologic hammer and sterile chisel, and rock samples, preserved in sterile plastic bags, transported, and stored at −20 °C in the Culture Collection of Antarctic fungi of the Mycological Section of the Italian Antarctic National Museum (MNA-CCFEE), until downstream analysis.

### Study data

In total, the dataset included 191 metagenomes, of which 100 have been assembled as described in Albanese et al.^11^. The remaining metagenomes were generated, sequenced, and assembled as described below. The final metagenomic set represented 149,585,625 metagenomic contigs.

### DNA extraction, library preparation, and sequencing

Total community DNA was extracted from 1 g of crushed rocks using DNeasy PowerSoil Pro Kit (Qiagen, German), quality checked by electrophoresis using a 1.5% agarose gel and Nanodrop spectrophotometer (Thermofisher, USA) and quantified using the Qubit dsDNA HS Assay Kit (Life Technologies, USA) according to Coleine et al.^10^. Shotgun metagenomic sequencing paired-end libraries were constructed and sequenced as 2×150 bp using the Illumina NovaSeq platform (Illumina Inc, San Diego, CA) at the Edmund Mach Foundation (San Michele all’Adige, Italy) and at the DOE Joint Genome Institute (JGI).

### Sequencing reads preparation and assembly

The metashot/mag-illumina v2.0.0 workflow (https://github.com/metashot/mag-illumina, parameters: --metaspades_k 21,33,55,77,99) was used to perform raw reads quality trimming and filtering, assembly and contigs binning on the metagenomic samples. In brief, adapter trimming, contaminant (artifacts and and spike-ins) and quality filtering were performed using BBDuk (BBMap/BBTools v38.79, https://sourceforge.net/projects/bbmap/). During the quality filtering procedure i) raw reads were quality-trimmed to Q6 using the Phred algorithm; ii) reads that contained 4 or more “N” bases, had an average quality below 10, shorter than 50 bp or under 50% of the original length were removed. Samples were then assembled individually with SPAdes v3.15.1^44^ (parameters –meta -k 21,33,55,77,99).

### Identification and clustering of viral genomes

Using a workflow similar to Guo et al.^45^, viral sequences were identified in metagenomic assemblies using VirSorter2 v. 2.2.3^25^ using --min-length 5000, --min-score 0.5, and --include-groups dsDNAphage,NCLDV,RNA,ssDNA,lavidaviridae. CheckV v0.8.1^31^ was run on the VirSorter2 predicted viral sequences using the “end_to_end” workflow VirSorter2 was then run again on the viral sequences from CheckV workflow with the --prep-for-dramv option. DRAM-v v. 1.2.2^40^ was then used to “annotate” sequences and then “distill” annotations into predicted auxiliary metabolic genes (AMGs) for phage.

Viral sequences were clustered into 95% similarity viral operational taxonomic units (vOTUs) using CD-HIT v. 4.8.1^26^ with the following parameters: -c 0.95 -aS 0.85 -M 0 -d 0. Prodigal v. 2.6.3^46^ was used to predict open reading frames in vOTUs using the -p meta option. VContact2 v. 0.9.19 was then run on predicted proteins from phage vOTUs and predicted proteins from the INPHARED August 2022 viral reference database to generate viral clusters (VCs) based on gene-sharing networks^27,28^. We assigned taxonomy to phage vOTUs based on VC membership as in Santos-Medellin et al^42^. Predicted viral sequences and 95% similarity vOTUS are archived on Zenodo^47^.

### Viral host-prediction

Hosts were predicted for the phage sequences identified using (i) a database of complete genomes from NCBI RefSeq, and (ii) a previously published database of representative metagenome-assembled genomes (MAGs) from Antarctic endolith samples. To produce (i), we used “ncbi-genome-download” to download all complete bacterial (n = 25,984) and archaeal (n = 416) genomes, as of April 7, 2022, from NCBI RefSeq^48^. For (ii), we downloaded MAGs from Zenodo (DOI: 10.5281/zenodo.7313591). We then used NCBI BLAST 2.12.0+ to convert these two databases into blast databases using “makeblastdb” and used “blastn” to compare vOTUs to these databases^34^. We filtered the blastn results in R based on existing thresholds^35,36,49^. Briefly, database matches had to share ≥ 2000 bp region with ≥ 70% sequence identity to the viral sequence and needed to have a bit score of ≥ 50 and minimum e-value of 0.001. Further to ensure matches did not represent partial or entirely viral contigs when searching against the MAG database, matches had to cover < 50% of the total MAG sequence length. As in Korthari et al.^36^, only the top five hits matching these thresholds were considered, with host predictions made at each taxonomic level only if the taxonomy of all hits were in agreement. Discrepancies resulted in no host prediction for that taxonomic level. We then combined host predictions from both the RefSeq and MAG databases together; if there were discrepancies between the two databases, we defaulted to the MAG-based prediction.

### Ecological analysis of vOTUs

We mapped reads from each metagenome to vOTUs using BBMap with a minid=0.90 to quantify vOTU relative abundance^50^. We then used SAMtools to convert resulting sam files to bam files and genomecov from BEDTools to obtain coverage information for each vOTU across each metagenome^51,52^. We then used bamM to parse bam files and calculate the trimmed pileup coverage (tpmean), which we used here in our analysis of viral relative abundance ^53^. We removed vOTUs which displayed < 75% coverage over the length of the viral sequence and viral sequences < 10 kbp in length prior to downstream analyses in R^54^. Thresholds for analysis of vOTUs were based on community guidelines for length (i.e. ≥ 10 kbp), similarity (i.e. ≥ 95% similarity), and detection (i.e. ≥ 75% of the viral genome length covered ≥ 1x by reads at ≥ 90% average nucleotide identity)^29,30^. To be conservative, we also removed vOTUs with a CheckV quality score of “not-determined” prior to downstream analysis. The viral abundance (tpmean), quality, taxonomy and annotation results were imported, analyzed, and visualized in R using many packages including tidyverse and phyloseq^55,56^. Analysis scripts associated with this study are on GitHub and archived in Zenodo^57^.

To compare viral diversity between metagenomes (i.e., beta diversity), we calculated the Hellinger distance, the Euclidean distance of Hellinger transformed abundance data. We performed Hellinger transformations using the transform function in the microbiome R package, calculated the Hellinger distance using the ordinate function in phyloseq, and then visualized these distances using principal-coordinate analysis (PCoA). We performed permutational multivariate analyses of variance (PERMANOVAs) with 9,999 permutations to test for significant differences in mean centroids using the model: Distance ∼ Site + Rock type. Models were tested with “by = margins” and “by = terms” with all sequential combinations. We ran the ordistep and ordiR2step functions to help assess optimal parameters to include in the model. Since PERMANOVA tests are sensitive to differences in group dispersion, we also tested for significant differences in mean dispersions using the betadisper and permutest functions from the vegan package in R with 9,999 permutations.

To test for correlations between viral community distances (Hellinger distances) and geographic distances, we first subset the data to exclude metagenomes from the Antarctic Peninsula, and to account for variation between rock types, subset the data to include only metagenomes representing sandstone samples. We calculated geographical distances between metagenomes using the distm function in the geosphere package in R. We performed Mantel tests in the vegan R package to assess correlations between the community and geographic distances using 9,999 permutations. Mantel tests were repeated with exclusion of community distances when the geographic distance was zero to assess if patterns persisted in the absence of data from the same site.

## Data availability

Metagenomes raw data are available under the NCBI accession numbers listed in Supplementary Table 1. Analysis scripts and intermediate data files associated with this study are on GitHub (https://github.com/stajichlab/Antarctic_Virus_Discovery) and archived in Zenodo (https://doi.org/10.5281/zenodo.7374327). Fasta files representing the entire catalog of predicted viral sequences and 95% similarity vOTUS are archived and available on Zenodo (https://doi.org/10.5281/zenodo.7245811).

## Acknowledgements

C.L.E. is supported by the National Science Foundation (NSF) under a NSF Ocean Sciences Postdoctoral Fellowship (Award No. 2205744). M.S. was supported by the NSF REU The National Summer Undergraduate Research Project, Award No. 2149582. C.C. is supported by the European Commission under the Marie Sklodowska-Curie Grant Agreement No. 702057 (DRYLIFE). C.C. and L.S. wish to thank the Italian National Antarctic Research Program for funding sampling campaigns and research activities in Italy in the frame of PNRA projects. The Italian Antarctic National Museum (MNA) is kindly acknowledged for financial support to the Mycological Section of the MNA and for providing rock samples used in this study stored in the Culture Collection of Antarctic fungi (MNA-CCFEE), University of Tuscia, Italy. M.D-B. is supported by a project from the Spanish Ministry of Science and Innovation (PID2020-115813RA-I00), and a project of the Fondo Europeo de Desarrollo Regional (FEDER) and the Consejería de Transformación Económica, Industria, Conocimiento y Universidades of the Junta de Andalucía (FEDER Andalucía 2014-2020 Objetivo temático ‘01 – Refuerzo de la investigación, el desarrollo tecnológico y la innovación’) associated with the research project P20_00879 (ANDABIOMA). J.E.S. is a CIFAR fellow in the Fungal Kingdom: Threats and Opportunities program. Data analyses performed at the High-Performance Computing Cluster at the University of California Riverside in the Institute of Integrative Genome Biology were supported by NSF grant DBI-1429826 and NIH grant S10-OD016290. Part of this work (proposals 10.46936/10.25585/60000791 and 10.46936/fics.proj.2020.51548/60000213) was conducted by the U.S. Department of Energy Joint Genome Institute (https://ror.org/04xm1d337), a DOE Office of Science User Facility, supported by the Office of Science of the US Department of Energy under Contract No. DE-AC02-05CH11231. The funders had no role in study design, data collection and interpretation, or the decision to submit the work for publication.

## Authors contribution

C.L.E., J.E.S., and C.C. conceived and designed the study. L.S. collected the samples. C.P, T.G.R. and S.T. oversaw and managed the metagenome sequencing and standard analysis. C.L.E. performed bioinformatic and statistical analysis with contributions from J.E.S. and M.S.. C.L.E. and C.C. interpreted results with contributions from M.S. and S.R.. C.L.E. and C.C. wrote the paper with contributions from all co-authors.

## Notes

https://doi.org/10.5281/zenodo.7374327

https://doi.org/10.5281/zenodo.7245811

### Competing Interest Statement

The authors have declared no competing interest.

