## Supplemental Table Legends and Figures for "Highly diverse and unknown viruses may enhance Antarctic endoliths’ adaptability"

[e]*Laboratorio de Biodiversidad y Funcionamiento Ecosistémico. Instituto de Recursos Naturales y Agrobiología de Sevilla (IRNAS), CSIC, Av. Reina Mercedes 10, E-41012, Sevilla, Spain*

[f]*Unidad Asociada CSIC-UPO (BioFun). Universidad Pablo de Olavide, 41013 Sevilla, Spain*

[g]*Research and Innovation Centre, Fondazione Edmund Mach, Via E. Mach 1, 38098, San Michele all'Adige, Italy*

[h]*Department of Energy Joint Genome Institute, Lawrence Berkeley National Laboratory, One Cyclotron Road, Berkeley, CA, 94720, USA*

[i]*Institute for Integrative Genome Biology, University of California, Riverside, Riverside, CA, USA.*

### Supplementary Table Legends and Figures:

**Table S1. Sample information and viral identification statistics from Antarctic metagenomes.** Here we provide information for each Antarctic metagenome explored here including site name, geographic area, rock type, year of collection, latitude, longitude, and SRA Accession numbers. This table also reports for each metagenome the number of predicted viral sequences, the number of proviral sequences, the number of vOTUs, the number of VCs, the average viral sequence length, average number of genes per sequence and the number of viral sequences identified by CheckV as being Complete, High-quality, Medium-quality or Low-quality. Only the viral sequences in this study that met the set thresholds for inclusion based on length (i.e. ≥ 10 kbp), similarity (i.e. ≥ 95% similarity), detection (i.e. ≥ 75% of the viral genome length covered ≥ 1x by reads at ≥ 90% average nucleotide identity), and quality (i.e., exclusion of viruses with "not-determined" CheckV scores) are summarized in this table.

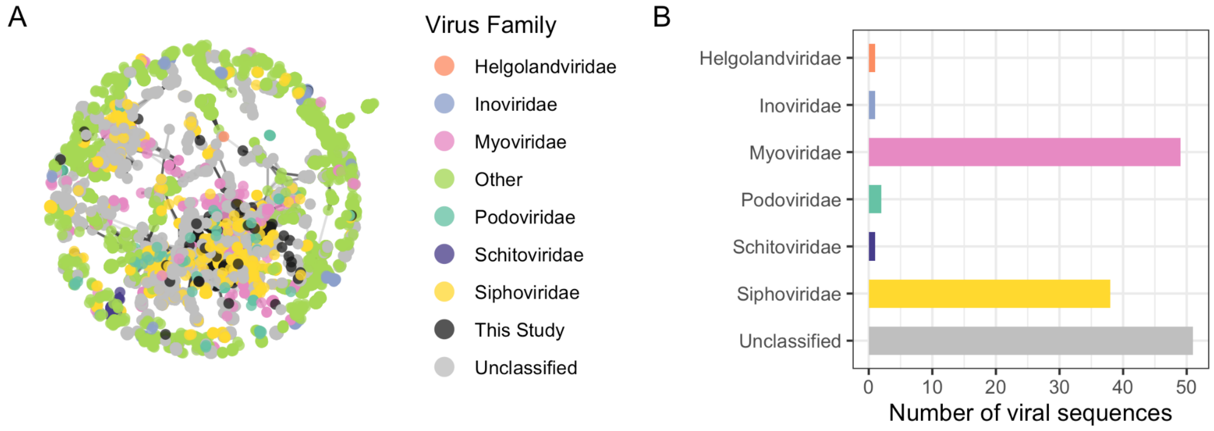


**Figure S1. Taxonomic classification of viral clusters (VCs) that include reference genomes.** (A) Subset of gene-sharing network from VContact2 showing only VCs that include reference genomes. Each node is a vOTU colored by predicted historical viral taxonomic family, with vOTUs identified in this study in black. Edges represent shared genes. (B) Bar chart displaying the number of viral sequences assigned to historical viral families based on VC membership with reference genomes.

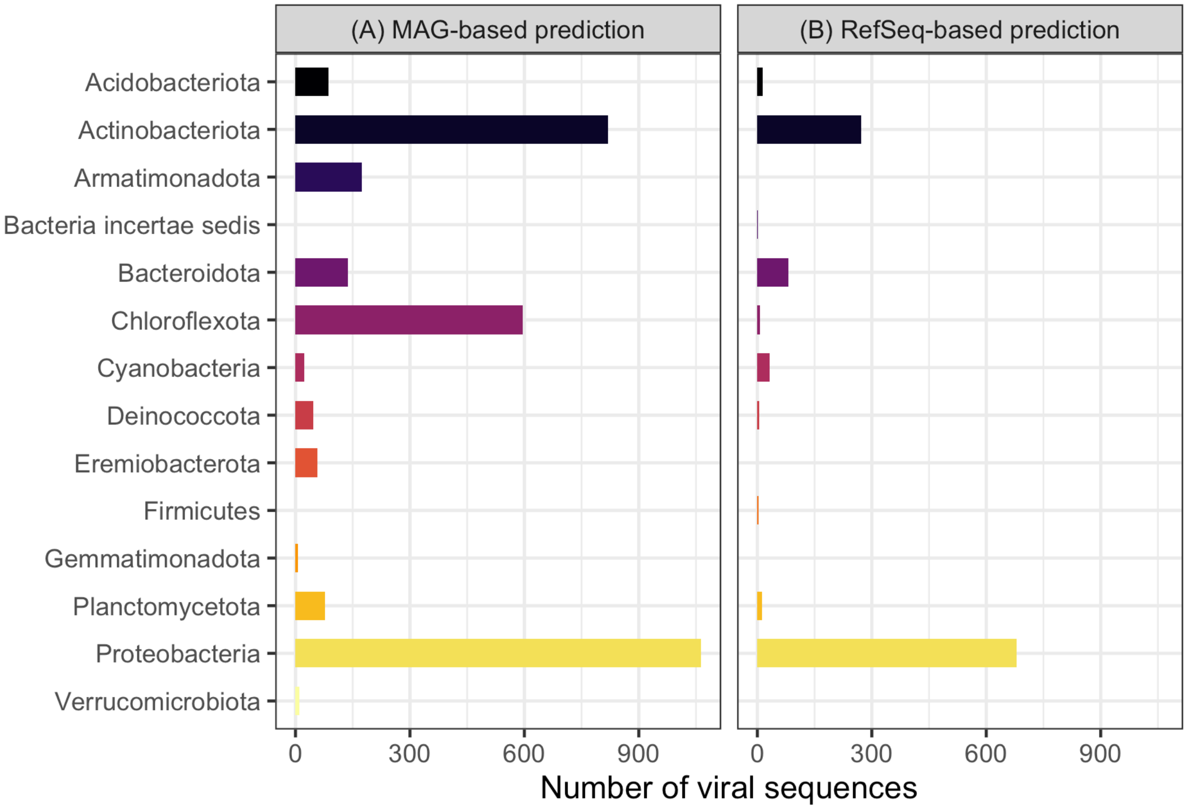


**Figure S2. Comparison of host predictions between MAG and RefSeq databases.** Bar charts displaying (A) MAG-based host predictions colored by predicted phylum, and (B) RefSeq-based host predictions colored by predicted phylum. Displayed are predictions for viral sequences that met the set thresholds for inclusion based on length, similarity, detection, and quality; viral sequences with no host prediction are not shown. Overall, 16.5% of viral sequences were assigned hosts based on MAG-based predictions, 3.0% were assigned hosts based on Refseq predictions, and 4.5% were assigned hosts based on both methods.
